# Abnormal spontaneous neural activity of auditory cortex and auditory pathway in tinnitus: a resting-state fMRI study

**DOI:** 10.1101/241216

**Authors:** Wei-Wei Cai, Jian-Gang Liang, Zhi-Hui Li, Yu-lin Huang, Li Wang, Tao Zhang

**Affiliations:** Department of Otolaryngology-Head & Neck Surgery, The First Affiliated Hospital of Jinan University, Guangzhou, Guangdong 510630, P.R. China; Department of Otolaryngology-Head & Neck surgery, Panyu Central Hospital, Guangzhou, Guangdong 511400, P.R. China; Department of Radiology, Guangzhou Panyu Central Hospital, Guangzhou, Guangdong 511400, P.R. China

**Keywords:** Resting-state functional magnetic resonance imaging, amplitude of low-frequency fluctuation, tinnitus, auditory cortex, auditory pathway, spontaneous neural synchrony

## Abstract

This resting-state functional magnetic resonance imaging (rs-fMRI) study in tinnitus patients was conducted to observe the spontaneous neural activity of the central auditory system using a derived index, mean amplitude of low-frequency fluctuation (mALFF). Tinnitus subjects with right-ear hearing impairment (THL) and without hearing loss (TNH) and two age-, sex-, and education-matched control groups (NC1 and NC2) were recruited for rs-fMRI. mALFF maps of the tinnitus and matched NC groups were plotted in the central auditory system, including the primary auditory cortex (PAC), higher auditory cortex (HAC), and hubs of the central auditory pathway. mALFF values of the activity clusters in the central auditory system of THL and TNH patients were extracted and correlated with each clinical characteristic. Significantly increased mALFF clusters were found in bilateral PAC and HAC of THL-NC1 maps and in the left inferior colliculus and right HAC of TNH-NC2 maps. Thus, subgroups of tinnitus with and without hearing impairment might exhibit different homeostatic plasticity in the central auditory system. mALFF values of aberrant active clusters in the central auditory system are partly associated with specific clinical tinnitus characteristics.

## Introduction

The contemporary view of the mechanism of tinnitus is that the absence of an input from the auditory periphery induces numerous plastic readjustments and aberrant states of activation, including hyperactivity, bursting discharges, and increases in neural synchrony in the central auditory and non-auditory systems (Mühlnickel, et al.,1998; Daniel A. Llano and Caspary,2012; Mulders and Robertson,2013; Chen, et al.,2014). The prevalence of tinnitus ranges from 10-15% in the adult population. (Xu, et al.,2011; Hoffman,2014). Hearing impairment, increasing age, and male sex have been identified as the most relevant risk factors for tinnitus (Hoffman,2014). Many patients with tinnitus have reported symptoms such as frustration, annoyance, irritability, anxiety, depression, hearing difficulties, hyperacousis, insomnia, and concentration difficulties. These symptoms are highly relevant to tinnitus severity and impair patients’ quality of life (Langguth,2011).

An important aspect during the onset of tinnitus is the spontaneous neural active changes in the central auditory system. Evidence from animal models of tinnitus indicates that abnormal states of neuronal activity in the auditory cortex and auditory pathway are the source of tinnitus in humans (Dong, et al.,2010; Ralli, et al.,2010). However, the main disadvantage of using animal physiological models is that these have not directly proven that animals are experiencing tinnitus. More recently, methods have been developed to measure changes in hemodynamics in tinnitus patients, using functional magnetic resonance imaging (fMRI); these methods include measuring fMRI neural activation with sound stimulation in auditory regions (Smits, et al.,2007; Lanting, et al.,2008; Gu, et al.,2010), resting state of the auditory region and functional connectivity with coherent networks (Maudoux, et al.,2012, 2012; Chen, et al.,2014, 2015), and combining functional connectivity using resting-state fMRI (rsfMRI) and structural MRI (Landgrebe, et al.,2009; Husain, et al.,2011; Seydell-Greenwald, et al.,2014). The main findings, when measuring fMRI activation with sound stimulation in tinnitus, are as follows: elevated neural activity, upon sound stimulus, in the inferior colliculus and primary auditory cortex is related with hyperacousis but not tinnitus (Gu, et al.,2010). In studies of tinnitus using rsfMRI, there are three popular methods of analysis: seeding, graphic connectivity analysis, and independent component analysis (ICA). The main findings in these studies show abnormal relationships in tinnitus patients between the auditory resting-state network and the following networks: default-mode, limbic, dorsal attention, and visual networks (Burton, et al.,2012; Kim, et al.,2012; Maudoux, et al.,2012, 2012). The foundation of functional connectivity was a signal change in different cerebrum regions with the connectivity of structures in different brain regions. The following studies combined anatomical and functional connectivity within the same framework, using diffusion tensor imaging and blood oxygen level-dependent (BOLD) functional connectivity and reported altered connecting regions from the inferior colliculus and amygdala to the auditory cortex. The findings of these studies imply a mechanism for accompanying syndromes in tinnitus patients.

rs-fMRI was developed to provide a new method for evaluation of regional neural spontaneous activity level in low frequency (0.01-0.08 Hz) fluctuation (ALFF) of the BOLD signal change (Biswal, et al.,1995; Kiviniemi, et al.,2000; Zang, et al.,2007). Combining electroneurophysiological recordings and fMRI, many studies have suggested that the LFFs of BOLD fMRI signals are closely related to spontaneous neuronal activities (Lu, et al.,2007; Mantini, et al.,2007; Shmuel and Leopold,2008; Rusiniak, et al.,2015; Shtark, et al.,2015). Notably, ALFF is reported to be higher in gray matter than in white matter. (Biswal, et al.,1995) Previous studies have confirmed the use of ALFF as an index to assess the regional synchronization of neural activity in multiple neuropsychiatric diseases, including cognitive dysfunction, chronic tinnitus and premenstrual syndrome (Kiviniemi, et al.,2000; Chen, et al.,2014; Huang, et al.,2017; Lei, et al.,2017). In the field of tinnitus research, existing studies have largely focused on aspects of abnormal functional connectivity between the auditory cortex and other non-auditory regions (Chen, et al.,2014). These reports suggest that the source of phantom tinnitus originates from abnormal spontaneous neural activity in the central auditory system in animal studies (Daniel A. Llano and Caspary,2012; Mulders and Robertson,2013; Chen, et al.,2014).

Our study was conducted to characterize the relationship between regional, abnormal, spontaneous neural activity in the central auditory system and mean ALFF (mALFF) in multiple levels of the central auditory system. We hypothesized that tinnitus patients may exhibit different values of mALFF within activity clusters of the central auditory system, which might thereby correlate with clinical presentation of tinnitus.

## Materials and methods

### Subjects

This study utilized data from four groups of subjects: tinnitus with hearing loss group (THL), normal control group 1 (NC1), tinnitus without hearing loss group (TNH), and normal control group 2 (NC2). The tinnitus groups were recruited from outpatient clinics at the Otorhinolaryngology Department of Guangzhou Panyu Central Hospital and from the First Affiliated Hospital of Jinan University, Guangzhou, China, from March 2015 to May 2017. NC groups 1 and 2 were matched in age, sex, and education to THL and TNH, respectively. Both THL and TNH groups comprised 12 patients with reported right-ear-tinnitus, tinnitus duration of >6 months, ages of 18-55 years, and right-handedness. Pure-tone audiometry was performed with a clinical audiometer using six octave frequencies (0.25, 0.5, 1, 2, 4, and 8 kHz). The audio thresholds of THL and TNH patients are shown in Fig. 1 and Fig. 2.

**Fig. 1.**
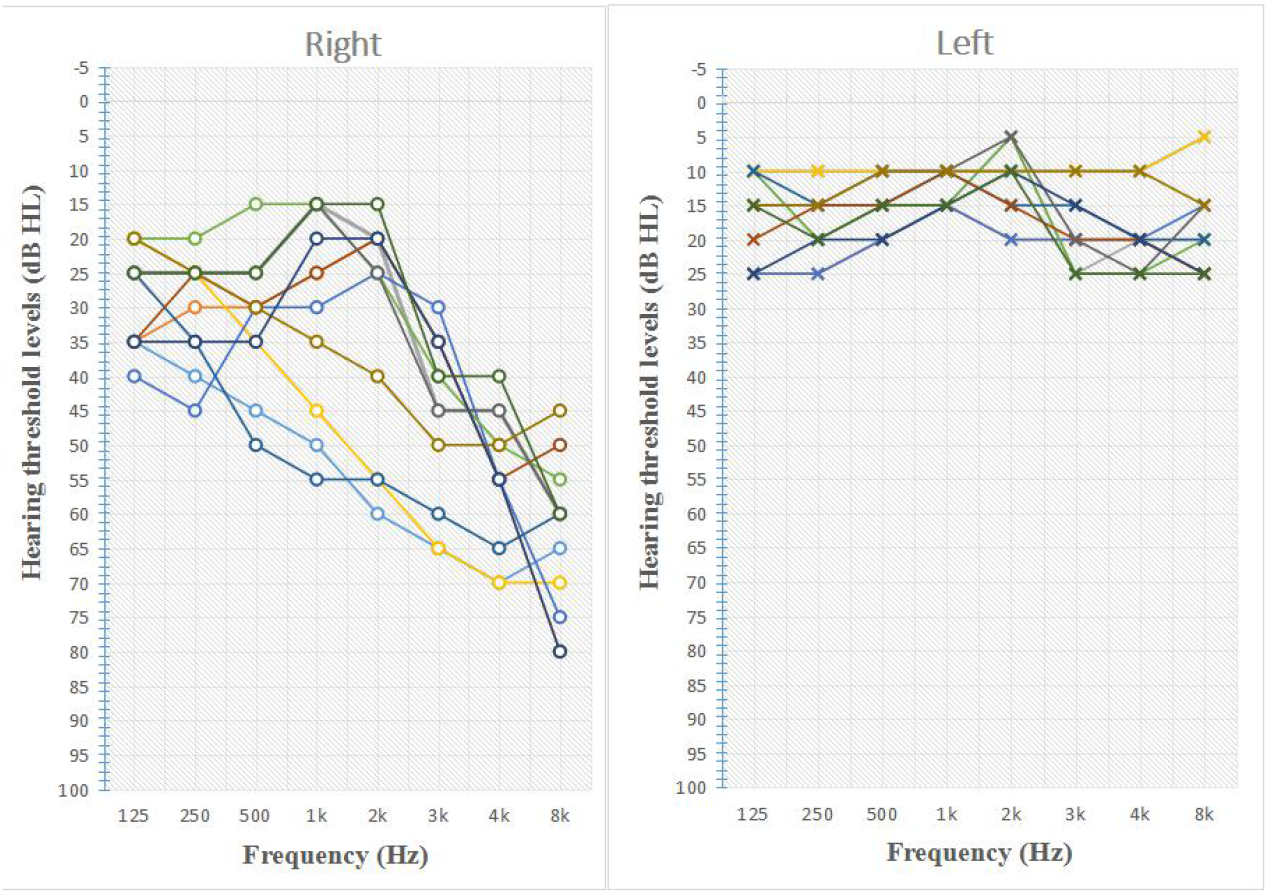
Characteristic hearing levels in the tinnitus with hearing loss (THL) group.

**Fig. 2.**
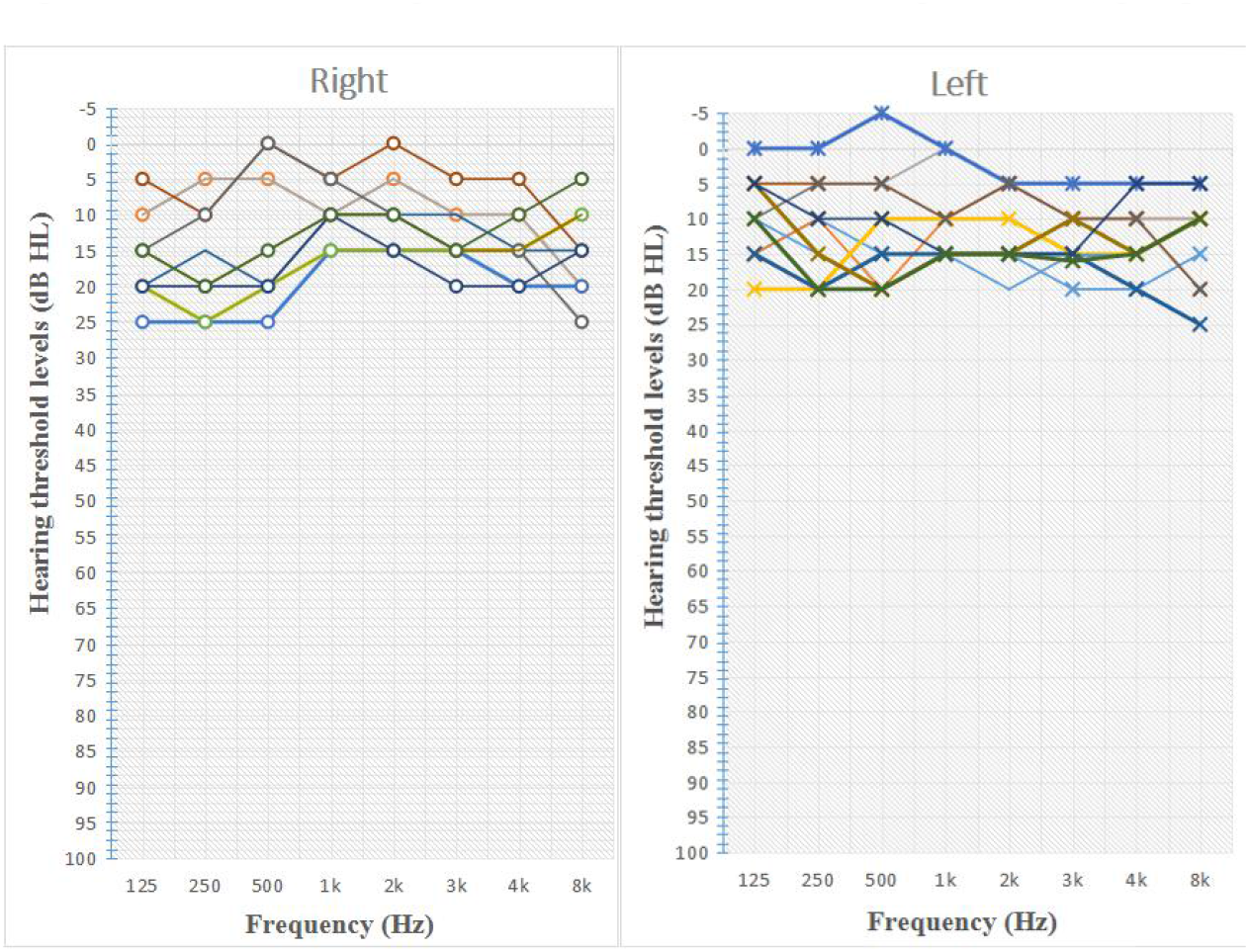
Characteristic hearing levels in the tinnitus without hearing loss (TNH) group.

The severity of tinnitus was assessed by a Chinese translation of the Tinnitus Handicap Inventory (THI), a self-reported tinnitus handicap questionnaire (Meng, et al.,2012). Group NC1 contained 30 subjects; group NC2 contained 30 subjects. All subjects were confirmed to have a Self-Rating Depression Scale (SDS) score of <50 and a Self-Rating Anxiety Scale (SAS) score of <50 (Zung,1971, 1986). All individuals provided written informed consent before participation in the study protocol. This study was approved by the Research Ethics Committee of Guangzhou Panyu Central Hospital and the First Affiliated Hospital of Jinan University, Guangzhou.

We excluded subjects with any of the following: Meniere disease; conductive deafness; alternative hearing level; cognitive or mental disorders; serious systemic diseases, such as heart failure or diabetes; epilepsy, alcoholism or use of psychiatric drugs; pregnancy; acoustic neuroma, brain stem or inferior colliculi diseases; hyperacousia; smoking; stroke; brain injury; Alzheimer’s disease; Parkinson’s disease. Table 1 summarizes the characteristics of tinnitus patients and NC subjects in this study.

### MRI data acquisition

All subjects underwent GE3.0T MRI (Discovery MR750) in the Imaging Center of the First Affiliated Hospital of Jinan University. Before MRI, all subjects lied in the examination bed with MRI noise-cancelling headphones and dual earplugs (Mack’s noise reduction ear plugs, USA) to reduce the noise level by 32 dB. To reduce head motion, we fixed a foam pad around the head of subjects. All subjects were asked to maintain a relaxed, resting state, including eyes closed, and to avoid serious thought, for approximately 20 minutes. BOLD signal acquisition conditions were defined as: 33 layers; repetition time (TR), 2000 ms; echo time (TE), 30 ms; layer thickness, 3 mm; flip angle, 90°; field of view (FOV), 200 x 200 mm; acquisition matrix, 64 x 64; and time points, 160. T1-weighted structure, using 3D T1-weighted fast gradient echo sequence, was performed as follows: sagittal scan, 175 layers of sagittal anatomical images; TR, 7.63 ms; TE, 3.74 ms; layer thickness, 1 mm; flip angle, 8°; FOV, 256 x 256 mm; and acquisition matrix, 256 x 256. The scanning range included the whole brain.

### Regions of interest and mask setting

The auditory cortex was utilized from the SPM anatomy toolbox v1.8 (http://www.fz-juelich.de/inm/inm-1/DE/Forschung/_docs/SPMAnatomy-Toolbox/SPMAnatomyToolbox_node.htm), including the primary auditory cortex (combination of area Te 1.0, 1.1, and 1.2) (Morosan, et al.,2001) and higher auditory cortex (Morosan, et al.,2005; NeuroImage,2005). The auditory pathway was based on Regions of Interest (ROI) reported by Muhlau et al. The ROI were were defined as spheres with a radius of 5 mm centered at ±10, -38, -45 (MNI coordinates) for the cochlear nuclei; ±13, -35, and -41 for the superior olivary complex; ±6, -33, -11 for the inferior colliculus; and ±17, -24, and -2 for the medial geniculate (Mühlau, et al.,2006). We define primary auditory cortex, higher auditory cortex, and all points in the auditory pathway, as the central auditory system mask, which we marked by Marsbar, a toolbox of region of interest analysis (Brett, et al.,2002).

### Data processing

All functional MRI data were processed by DPABI (a toolbox for Data Processing & Analysis for Brain Imaging) V2.3 (Yan, et al.,2016) for data processing, including: time correction, reorientation of T1 and functional image data, coregistration of T1 and functional image data, head motion correction, spatial normalization by diffeomorphic anatomical registration through exponentiated lie algebra, de-linear drift, filter processing (0.01-0.08 Hz), and cerebrospinal fluid signal processing. Gray matter images were extracted from T1 image data. ALFF results were divided by mean value of the whole brain to generate mALFF images.

### Statistical analysis

#### Statistical analysis in mALFF images between groups

We used the DPABI (a toolbox for Data Processing & Analysis for Brain Imaging, V2.3) (Yan, et al.,2016) statistical analysis function to perform calculations between groups of images. The two-sample *t*-test was used to compare mALLF maps of THL with NC1, and TNH with NC2, within the mask of the auditory system. Gray matter was added as a covariate.

We performed 5000 permutations tests using the Threshold Free Cluster Environment (TFCE) (Winkler, et al.,2016). The cluster forming threshold z was set to 3.1; a threshold-free cluster-enhanced technique (Nichols,2010) was used to control for multiple comparisons and evaluate the voxel-wise significant differences among the groups.

#### Correlation analysis

To investigate the relationship between the mALFF of activity clusters (in the central auditory system) and clinical data of tinnitus patients, we extracted mALFF values from activity clusters in the central auditory system of THL and TNH subjects. Subsequently, Pearson’s correlation coefficients were used to compare values of mALFF with each clinical characteristic of tinnitus subjects, via analysis by the corrplot tool kit of R software (Core,2015).

## Results

### Comparison clinical data between groups

There were no differences in clinical data between any tinnitus patient groups and any healthy control groups. (Table 1).

**Table 1.** Clinical characteristics of study subjects.

| **Table 1** |  | **Clinical characteristics of study subjects** |  |  |  |  |
| --- | --- | --- | --- | --- | --- | --- |
|  | THL | NC1 | *P1* value | TNH | NC2 | *P2* value |
| Gender (male/female) | 8/4 | 13/17 | 0.237 | 4/8 | 12/18 | 0.689 |
| Age (years) | 39.5±11.49 | 37.00±11.75 | 0.603 | 35.33±10.70 | 35±10.10 | 0.720 |
| Education (years) | 14±1.48 | 14.25±1.55 | 0.689 | 13.50±1.57 | 13.25±1.75 | 0.698 |
| SDS score | 38.67±9.36 | 36.25±8.04 | 0.661 | 34.67±7.30 | 35.25±6.33 | 0.274 |
| SAS score | 35.17±10.50 | 38.75±6.26 | 0.341 | 35.67±9.37 | 37.65±7.23 | 0.240 |
| Tinnitus ear<br>(Left/Right/Bilateral) | 0/13/0 | \ | \ | 0/10/2※ | \ | \ |
| THI score | 68.50±4.36 | \ | \ | 55.33±11.03 | \ | \ |
| Tinnitus duration (months) | 31.58±21.94 | \ | \ | 36.58±18.03 | \ | \ |
| Frequency range of tinnitus | 4–6 kHz | \ | \ | 5–8kHz | \ | \ |

※Two subjects have dominating right ear tinnitus. THL, tinnitus with hearing loss; NC1, normal controls group 1; TNH, tinnitus without hearing loss; NC2, normal controls group 2; SDS, Self-rating Depression Scale; SAS, Self-rating Anxiety Scale; THI, Tinnitus Handicap Inventory.

### Comparison of mALFF in auditory cortex and auditory pathway between groups

#### THL and NC1

Analyses, using two-sample *t*-tests, revealed significant clusters of mALFF maps between THL and NC1 subjects. Significantly increasing mALFF clusters can be found in bilateral higher auditory cortex (HAC, Fig. 3 A) and primary auditory cortex (PAC, Fig. 3 B). There were no significant different clusters in the auditory pathway (AP). Cluster characteristics are shown in Table 2.

**Fig. 3.**
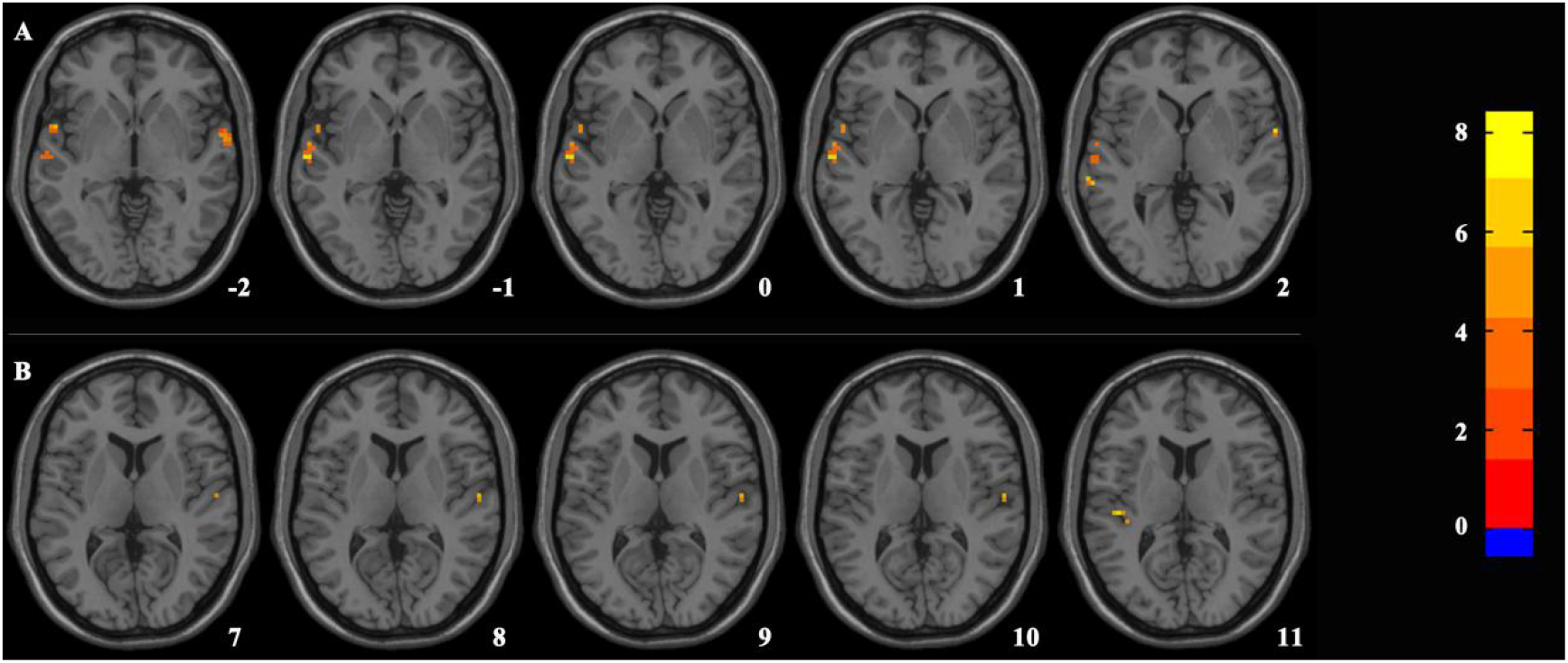
Significant mean amplitude of low-frequency fluctuation (mALFF) differences in tinnitus patients with hearing loss, compared with healthy control group 1. A shows different clusters of mALFF in higher auditory cortex. B shows different clusters of mALFF in primary auditory cortex. The 5000 permutations test was performed and the cluster-forming threshold was set to *p* = 0.001 (z = 3.1), with a threshold-free cluster-enhanced technique.

**Table 2.** Activity clusters of mean amplitude of low-frequency fluctuation (mALFF) of tinnitus patient groups, compared with healthy control groups, in auditory regions.

|  | Location | Number of | Sum Clusters size | Peak score | Peak MNI coordinates | mALFF value of active clusters |  |  |
| --- | --- | --- | --- | --- | --- | --- | --- | --- |
|  |  | clusters | Voxels (Left/Right) |  | x, y, z (mm) | TG | / | NC |
| THL | HAC | 2 | 49/29 | 8.501 | -63, -15, 0 | $1.871 \pm 0.185 / 1.054 \pm 0.089$ | | |
| | PAC | 2 | 4/3 | 7.245 | -45, -24, 12 | $1.873 \pm 0.119 / 1.173 \pm 0.242$ | | |
|  | AP | 0 | \ | \ | \ | \ |  |  |
| TNH | HAC | 1 | 0/64 | -4.433 | 69, -9, 6 | $0.830 \pm 0.126 / 1.086 \pm 0.145$ | | |
| | | 1 | 0/4 | 5.969 | 57, 3, -6 | $1.999 \pm 0.317 / 0.935 \pm 0.245$ | | |
|  | PAC | 0 | \ | \ | \ | \ |  |  |
| | AP* | 1 | 4/0 | 6.979 | -9, -33, -15 | $2.208 \pm 0.301 / 1.525 \pm 0.524$ | | |

AP\*: Auditory pathway, the peak MNI coordinates of active cluster was -9, -33, -15; this was located in left inferior colliculus, one regions of interest in auditory pathway, at -6, -33, -11; radius = 5 mm. THL, tinnitus with hearing loss; TNH, tinnitus without hearing loss; HAC, higher auditory cortex; PAC, primary auditory cortex. TG: tinnitus groups; NC: normal control

#### TNH and NC2

Analyses using a two-sample *t*-test revealed significant clusters of mALFF maps between TNH and NC2. Significantly different mALFF clusters were found in left PAC, left HAC(Fig. 4 A), and right AP (Fig. 4 B). Cluster characteristics are shown in Table 2.

**Fig. 4.**
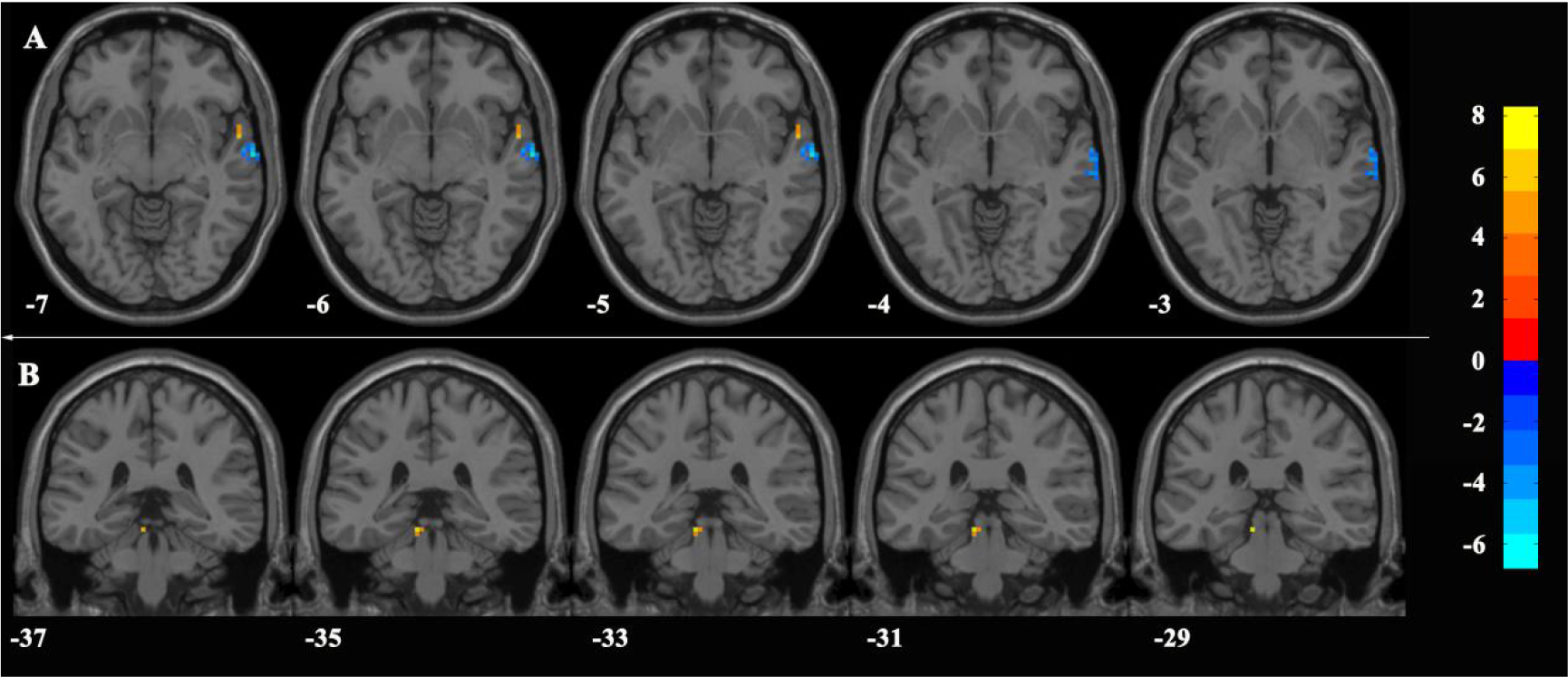
Significant mean amplitude of low-frequency fluctuation (mALFF) differences in tinnitus patients without hearing loss, compared with healthy control group 2. A shows different clusters of mALFF in higher auditory cortex. B shows different clusters of mALFF in the auditory pathway. The 5000 permutations test was performed and the cluster-forming threshold was set at *p* = 0.001 (z = 3.1), with a threshold-free cluster-enhanced technique.

### Correlation analysis between mALFF values of activity clusters in auditory system and tinnitus clinical data

In the THL group, the average mALFF values of activity clusters in HAC and PAC were extracted and correlated with the right ear average hearing threshold (RAHT), duration of tinnitus months (DTM), THI, SDS, and SAS. Color coding (Fig. 5) indicates the value of the coefficient of the Pearson correlations (R) between the mALFF values of activity clusters in the auditory cortex and clinical data. mALFF value correlated with the RAHT in PAC and HAC, THI in HAC, SDS in HAC, and SAS in HAC (correlation intensity: *p<0.01)* (Figs.6).

**Fig. 5.**
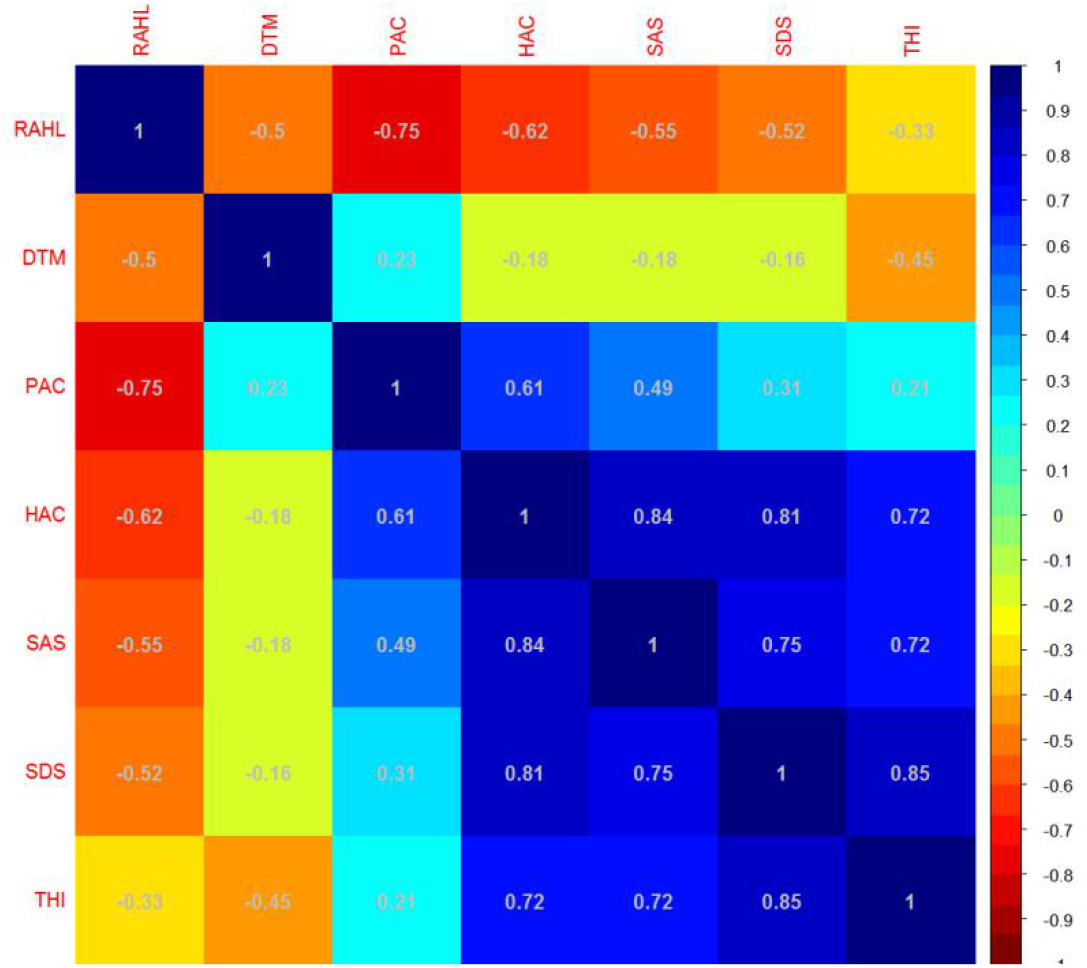
Color-coding indicates the value of the coefficient of Pearson correlations (R) between the mean amplitude of low-frequency fluctuation (mALFF) values of activity clusters in auditory cortex and clinical data. RAHT: Right-ear average hearing threshold; DTM: duration of tinnitus months; HC: value of activity clusters in higher auditory cortex. AC: mean amplitude of low-frequency fluctuation (mALFF) value of activity clusters in primary auditory cortex. DTM: duration of tinnitus months. SDS: Self-Rating Depression Scale. SAS: Self-Rating Anxiety Scale. THI: Tinnitus Handicap Inventory.

**Fig. 6.**
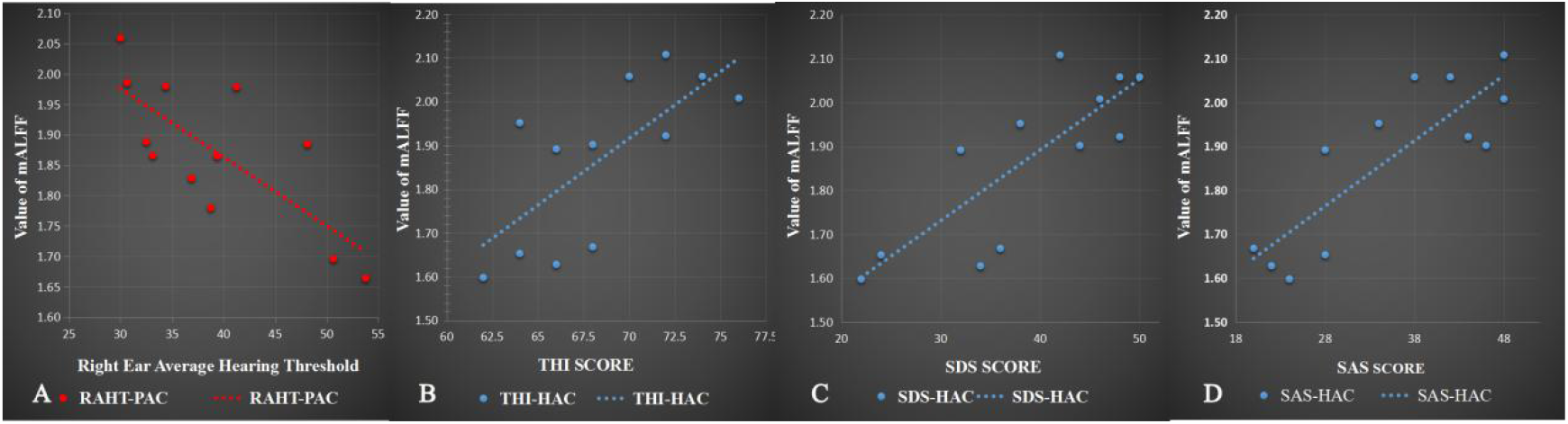
Pearson correlations between clinical parameters (RAHT, THI score, SDS score, and SAS score) and mean amplitude of low-frequency fluctuation (mALFF) values of activity cluster in primary auditory cortex and higher auditory cortex. (A) RAHL-PAC: r = -0.754, *p* = 0.005. (B) THI-HAC: r = 0.718, *p* = 0.009. (C) SDS-HAC: r = 0.814,*p* = 0.001. (D) SAS-HAC: r = 0.843,*p* = 0.001.

In the TNH group, the average mALFF values of activity clusters in HAC and AP were extracted and correlated with clinical data, including DTM, THI, SDS, and SAS. The color-coding (Fig. 7) indicates the value of the coefficient of Pearson correlations (R) between clinical data and the mALFF value of activity clusters in the auditory cortex. mALFF values correlated with the DTM, THI, SDS, SAS, and with the mALFF value in the activity cluster in the ROI of inferior colliculus (IC; ±6, -33, -11, radius = 5 mm) and active clusters in right HAC. The correlation intensity of DTM-IC, THI-IC, THI-IHAC, SDS-IC, SDS-DHAC, and SAS-DHAC were *p<0.01,* whereas the correlation intensity of SAS-IC was *p*<0.05. (Fig. 8).

**Fig. 7.**
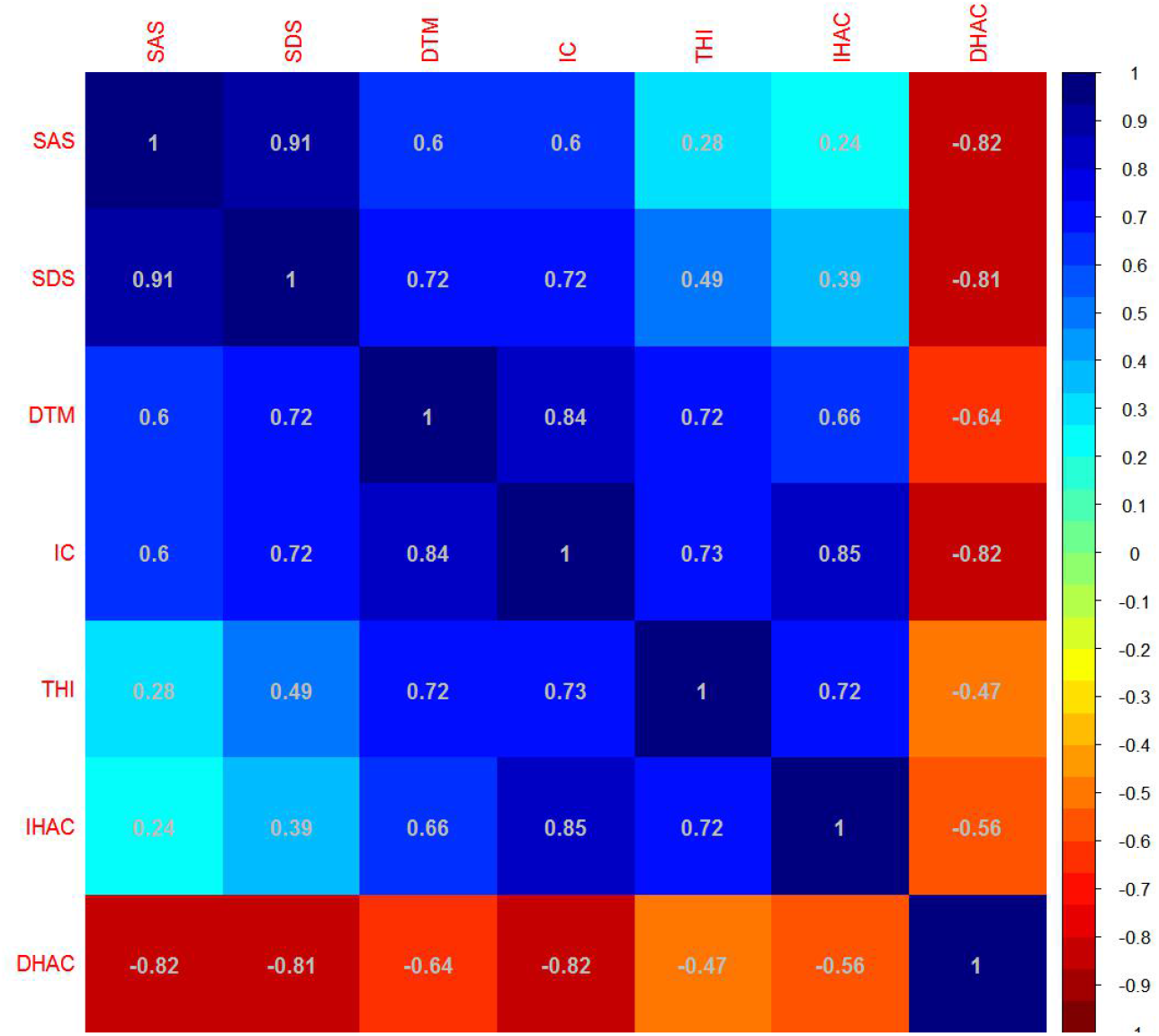
Color-coding indicates the value of the coefficient of Pearson correlation (R) between the mean amplitude of low-frequency fluctuation (mALFF) values of activity clusters in the central auditory system and clinical data. DTM: duration of tinnitus months, IC: value of mALFF in activity cluster of inferior colliculus. DTM: duration of tinnitus months. SDS Self-Rating Depression Scale. SAS: Self-Rating Anxiety Scale. THI: Tinnitus Handicap Inventory. IHAC: increased value of mALFF in active cluster of higher auditory cortex. DHAC: decreased value of mALFF in active cluster of higher auditory cortex.

**Fig. 8.**
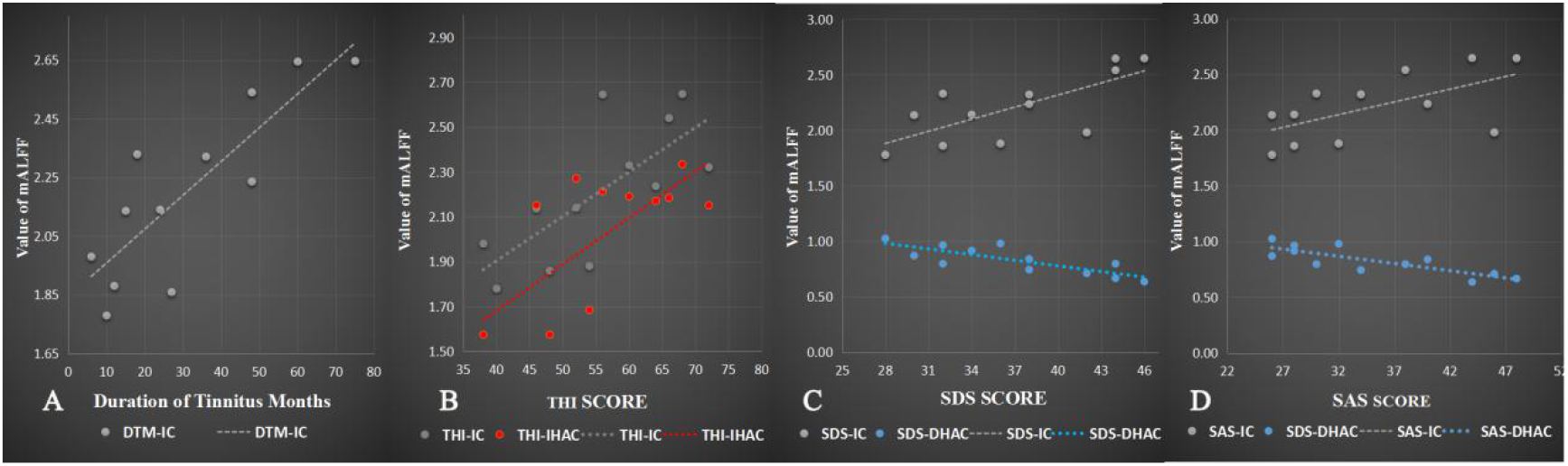
Pearson correlations between DTM, THI score, SDS score, SAS score, and mean amplitude of low-frequency fluctuation (mALFF) values of activity clusters in the HAC and left inferior colliculus (IC) (A) DTM-IC: r = 0.840,*p* = 0.001. (B) THI-IC, r = 0.734, *p* = 0.008. THI-IHAC: r = 0.722,*p* = 0.008. (C) SDS-IC, r = 0.723,*p* = 0.008. SDS-DHAC: r = -0.81,*p =* 0.001. (D) SAS-IC: r= 0.604,*p* = 0.038. SAS-DHAC: r= 0.820,*p* =

## Discussion

### Summary of results

In this study, we aimed to identify abnormal neural activity clusters in PAC, HAC, and AP of tinnitus patients, with and without hearing loss, using the mALFF index. In comparisons between THL and NC1, mALFF maps (Fig. 3) indicate increased values of mALFF in the clusters of bilateral PAC and HAC, but no differences in hubs of the AP, although all subjects in the THL group exhibited right-ear tinnitus with hearing impairment. The value of mALFF activity clusters in the PAC of THL was negatively correlated with RAHL and did not correlate with the THI score, as hyperacousis patients were excluded. The value of mALFF active clusters in HAC was positively correlated with THI, SAS, and SDS. In comparisons between TNH and NC2, mALFF maps (Fig. 4) reveal abnormal activity clusters in HAC, including both decreased and increased activity clusters, and increases in the IC. DTM, THI, SAS, and SDS were positively correlated with mALFF value of the activity cluster in IC.

### Why was mALFF used as an index to evaluate spontaneous neural baseline activation of the central auditory system?

Spontaneous neuron activity includes spontaneous firing rate, synchrony between neurons, and busting in specific neural structure, as physiological correlates of tinnitus (Kaltenbach,2011). Increased synchrony between neurons in the central auditory system could create sound perception (Langers, et al.,2007); thus, it is feasible that increased synchrony in the absence of peripheral auditory system could lead to the perception of a phantom sound (Noreña and Eggermont,2003).

The mechanism of the amplitude of low-frequency fluctuation (ALFF) of the rs-fMRI signal has been demonstrated by several studies, using comparison with epidural electroencephalography (EEG) (Lu, et al.,2007; Mantini, et al.,2007; Shmuel and Leopold,2008; Shtark, et al.,2015). At the level of neural ensembles, a synchronized activity of large numbers of neurons can lead to macroscopic oscillations, which can be observed through EEG and correlated with values of mALFF. In functional connections of rat primary somatosensory cortices, Lu et al. found that, unlike the evoked fMRI response, which correlates with power changes in high-frequency bands, the power coherence in low-frequency bands correlates with mALFF values in the BOLD signal on rs-fMRI, in a region-specific and dose-dependent fashion. (Lu, et al.,2007) Further, the ALFF of the BOLD signal on rs-fMRI correlates with power coherence in low-frequency bands, particularly the delta band of EEG (Lu, et al.,2007). Another study demonstrated that the theta-gamma band of neuronal activity in EEG correlates with spontaneous fluctuations in fMRI signals from the primary and secondary auditory cortex (Shmuel and Leopold,2008; De Ridder D, et al.,2011). In human brains studied by rs-fMRI, each of these networks, deprived by independent components analysis, could be generally associated with more than one kind of electric oscillation, confirming that the brain does not generally display pure rhythms within distinct frequency bands, mostly in restricted neuronal circuits, but shows a coalescence of rhythms, as demonstrated by maps obtained with the previous EEG/fMRI studies on alpha-power correlation in the auditory-phonological system (Mantini, et al.,2007). To summarize, the main findings of previous studies about correlations between ALFF and EEG were as follows. First, ALFF is an index that can be used to evaluate the level of spontaneous neural synchrony in the resting-state, but not in the evoked brain. Second, ALFF of the BOLD signal on rs-fMRI in human brain correlates with the low frequency band fluctuation, (i.e, alpha, delta, gamma, and theta bands on EEG), which reflects synchronized activity of large numbers of neurons that can lead to macroscopic oscillations and can be observed by ALFF of rs-fMRI. Finally, the brain typically does not display pure rhythms within distinct frequency bands, but exhibits a coalescence of rhythms (Mukamel, et al.,2005). ALFF has been used as an index to evaluate spontaneous neural synchrony in prior studies (Biswal, et al.,1995; Kiviniemi, et al.,2000; Chen, et al.,2014; Lei, et al.,2017). Thus, mALFF was used as an index in our study to evaluate the spontaneous neural synchronization of the central auditory system.

### Abnormal spontaneous neural activity of central auditory cortex in previous studies of tinnitus with and without hearing loss

#### Tinnitus with hearing impairment

Most studies have suggested that the mechanism of tinnitus with hearing loss, absent input from the auditory periphery as a trigger, induces aberrant states of neural activation and reorganization in the central auditory system. (Daniel A. Llano and Caspary,2012; Mulders and Robertson,2013; Chen, et al.,2014) Studies of animal and human images indicate that hearing loss and tinnitus are factors that contribute to elevation in the baseline neural activity of PAC, which induces tinnitus perception. (Engineer, et al.,2011; Yang, et al.,2011; Daniel A. Llano and Caspary,2012; Mulders and Robertson,2013; Chen, et al.,2014) In human research, Peyman et al. combined EEG with MRI to observe patients suffering tinnitus with hearing loss (Peyman, et al.,2012), and found that tinnitus with hearing loss exhibited significant enhancement in the power of the delta band wave in auditory cortex, which was reduced after tinnitus masking. Other studies have similarly suggested that baseline neural activity, as detected by EEG, in PAC can be altered by tinnitus inhibition. (van der Loo E, et al.,2009; Roberts, et al.,2015) This implies that the baseline of spontaneous neural synchrony in PAC can be affected by both hearing threshold and tinnitus. (Kahlbrock and Weisz,2008; van der Loo E, et al.,2009; Peyman, et al.,2012) Studies of tinnitus with hearing loss using auditory steady-state response (ASSR), magnetoencephalography (MEG), and sound-evoked fMRI suggest that abnormal baseline neural activity in PAC correlates with severity of tinnitus (Diesch, et al.,2010, 2010) and hearing threshold. (Gu, et al.,2010) In our study, we found that abnormal hyper-neural activity clusters in PAC correlate with average hearing threshold but not with THI score, suggesting that mALFF of activity clusters in PAC can detect a coalescence of rhythms of spontaneous neural synchronic activity (dominated by input deficit), but cannot observe differences between THL and NC1 for individual tinnitus pitches (as in the THL group).

In most mechanistic studies of tinnitus with hearing loss, researchers are primarily concerned with abnormal spontaneous neural synchrony in PAC for direct induction of tinnitus perception. Thus, the function of HAC seems to be neglected in tinnitus research. In our study, we found that mALFF value in activity clusters in HAC of the THL group was correlated with THI, as well as SAS and SDS. Moreover, SAS and SDS scores were more intensely correlated with mALFF values of the HAC than THI score, implying that abnormal neural activity of HAC affects the severity of tinnitus, but not tinnitus perception, in THL patients.

The function of HAC was investigated in some studies, suggesting that aberrant neuronal activity in the HAC correlates with selective attention for sound perception, emotional state, and other cognitive processes (Plichta, et al.,2011; Brosch, et al.,2013; Popovych and Tass,2014; Seither-Preisler, et al.,2014; Eggermont and Tass,2015; Balconi and Vanutelli,2016). Thus, the baseline neural activity in HAC might indirectly affect the severity of tinnitus in patients. Paul et al. found that HAC exhibited an increased amplitude, as measured by ASSR, in the control group, but not in the tinnitus with hearing loss group, during tests where both groups were requested to give their attention to the contents of sound stimulation (Paul, et al.,2014), suggesting that an enhancement of baseline neural activity in HAC of the THL group might reduce the ability to react to sound stimulation (during the ASSR assessment) and induce dysfunctional selective attention. An early fMRI study suggests that chronic electrical stimulation of the HAC can suppress tinnitus-based depression score, but does not affect the loudness of tinnitus (Friedland, et al.,2007). Similarly, spontaneous neural activity of HAC, as detected by fMRI, in tinnitus patients with hearing loss was downgraded after effective treatment, and a sub-score of the tinnitus functional index—the relaxation score—significantly decreased (Emmert, et al.,2017). Furthermore, it follows that amplitude of the ASSR was modulated after residual inhibition in PAC, but not in HAC (Roberts, et al.,2015)

Spontaneous neural synchronic modulation in PAC, as evaluated by mALFF, might not correlate with perception of tinnitus, but does correlate with average hearing threshold; in contrast, change in the HAC affects sensory processes of tinnitus patients, including the dysfunction of selective attention, specifically emotional and cognitive processes, thus correlating with severity of tinnitus.

#### Tinnitus without hearing impairment

Although most tinnitus was induced by obvious hearing loss, some tinnitus patients without hearing loss might exhibit severe tinnitus (Savastano,2008; Martines, et al.,2010). Hidden hearing loss is suspected in this type of tinnitus, which differs from tinnitus with obvious hearing loss.

In our study, increased mALFF value of the active cluster occurred in contra-IC of the right-ear tinnitus patients within the normal audiogram group, implying that an elevated baseline of spontaneous neuronal synchrony occurred in the IC of normal hearing patients. Evidence of hyperactive IC has been found in the animal behavioral tinnitus model (Mulders and Robertson,2009, 2011). However, this hyperactivity may not be a sufficient sole generator of tinnitus for increased spontaneous neural activity, as it was present even in the absence of behavioral evidence of tinnitus (Coomber, et al.,2014; Ropp, et al.,2014). In auditory nerve fiber deafferentation studies of tinnitus with normal hearing audiograms, studies in humans have shown that the centrally generated wave V occurs as result of reduction in wave I auditory brainstem response (ABR) amplitude, suggesting that the auditory pathway of middle brain might exhibit homeostatic plasticity to elevated neuronal response gain (Aitkin and Phillips,1984; Schaette and McAlpine,2011; Hesse, et al.,2016). Tinnitus sound-evoked fMRI also found that IC showed increased activity after sound stimulation (Smits, et al.,2007; Lanting, et al.,2008; Gu, et al.,2010), but these studies did not exclude hyperacousis (Gu, et al.,2010). Additionally, clinical tinnitus studies offer indirect evidence that modulated somatosensory input affects neural activity of the IC and correlates with the onset of tinnitus (Lanting, et al.,2010; Gritsenko, et al.,2014). Our finding is consistent with the accumulated evidence for hyperactive IC in tinnitus.

The IC is a near-obligatory relay in the ascending auditory pathway, a point where virtually all lemniscal and extra-lemniscal ascending inputs converge(Aitkin and Phillips,1984; Marsh, et al.,2002; Bajo and Moore,2005) and is no longer considered solely as an organizing center for integration of auditory processing, but as a key organ in human emotion response that utilizes considerable non-auditory input (Marsh, et al.,2002; Bajo and Moore,2005; Chen, et al.,2015; Kim, et al.,2017). In our study, the mALFF value of an abnormal neural active cluster in left IC is positively correlated with DTM, THI, SAS, and SDS, supporting previous evidence that modulated spontaneous neural activity of the IC affects emotion and tinnitus severity.

In our study, the mALFF value of the left IC was positively correlated with DTM. One possible etiology for tinnitus with hidden hearing loss is non-traumatic noise exposure, which suggests that the increased spontaneous activity of IC is dependent on external auditory input in the early stages(Coomber, et al.,2014; Ropp, et al.,2014) and can be modulated by ablation of the dorsal cochlear nucleus within 6 weeks of noise exposure(Mulders and Robertson,2009). The hyperactivity of the IC subsequently becomes self-sustaining, as result of homeostatic plasticity in the IC, which is induced by 3-5 months of acoustic exposure(Brozoski and Bauer,2005). However, none of these studies offered a correlation between tinnitus duration and the level of spontaneous neural activity of the IC.

The IC has been characterized as a major locus for the integration of excitatory and inhibitory inputs (Eggermont,2001). As a result of homeostatic plasticity in the IC, the gain to supplement input deficiency at hearing loss frequencies might accompany neural noise and cortical tonotopic map distortion, which would induce the perception of tinnitus(Robertson and Mulders,2012). In our study, we did not find hyperactive clusters in the PAC of the mALFF map for various conditions in the TNH group, including individual tinnitus pitch, different frequency ranges of hidden hearing impairment, and various environmental pathogenic factors. In HAC, we found decreased and increased activity clusters occur at the same time and that decreased mALFF values of the cluster in right HAC are strongly negatively correlated with the right IC (Fig. 7). In an early animal study, auditory neural fibers were found between IC and HAC in the cat (Eggermont and Kenmochi,1998). More recently, functional connections between the IC and auditory cortex have been reported in some studies (Bajo and Moore,2005; Chen, et al.,2015). Taken together, these prior findings and our results that the mutual interaction between IC and HAC might be the result of homeostatic plasticity in IC.

### Significance of the results

In our study, abnormal spontaneous neural activity in the auditory system showed that THL and TNH might exhibit different homeostatic plasticity because of their location in the central auditory system. These subgroups of tinnitus exhibit different spontaneous neural activity in different regions of the auditory system.

These findings suggest that different types of tinnitus can be evaluated by different regions of the auditory system and should be managed by different treatments, such as repetitive transcranial magnetic stimulation(De, et al.,2011), neurofeedback(Emmert, et al.,2017), and transcutaneous electrical nerve stimulation (Tyler, et al.,2017). In future studies, we will collect more mALFF data from the auditory cortex and auditory pathway of healthy subjects, and will evaluate relationships with age, years of education, and sex, to be used as a standard index to evaluate severity and classification of tinnitus.

### Study limitations

First, our study observed the relationship between an abnormal mALFF of the auditory system and tinnitus characteristics in a cross-sectional format with a small sample size (n < 30). Therefore, future studies must analyze a larger subgroup of tinnitus cases and include lateral tinnitus, hearing impairment grading, age grouping, and multiple etiologies, across multiple centers. Moreover, we cannot completely eliminate the substantial acoustic noise produced during the fMRI procedure with ear plugs and MRI noise-cancelling headphones, which might affect baseline neural activity (Rondinoni, et al.,2013). Finally, the neural activity of special regions in the auditory system might be modulated by attention in consciousness (Jäncke, et al.,1999; Martinez-Granados, et al.,2014; Ghazaleh, et al.,2017) and we cannot be sure that all subjects followed all requirements, including keeping eyes closed, staying relaxed, and avoiding any thinking.

## Conclusion

The subgroups of tinnitus with and without hearing impairment might exhibit different homeostatic plasticity at the level of the central auditory system; THL showed an abnormal spontaneous neural active cluster in HAC, correlating with tinnitus characteristics, whereas TNH showed abnormal activity in the IC. Thus, the value of mALFF in the auditory system partly reflects the severity of tinnitus in specific situations.

## Acknowledgments

The authors thank Prof. Xiang-Li Zeng, Department of Otolaryngology-Head & Neck Surgery, The Third Affiliated Hospital of Sun Yat-sen University, China. This study was supported by the Science and Technology Planning Project of Panyu District of GuangZhou City, China (Grant Number: 2015-Z03-41) and by the Medical Master and Doctor Foundation of the Central Hospital of Panyu District, Guangzhou, China (Grant Number: 2016-S-03).

